# PCV2 aggravated SS2 infection directly showed from differentially expressed proteome of PK15 cells

**DOI:** 10.1101/2020.02.24.962738

**Authors:** Tan Jimin, Xiao Qi, Li Xingguang, Fan Hongjie, He Kongwang

**Author notes:** Correspondence: Fan Hongjie, He Kongwang.

## Abstract

PCV2 and SS2 were clinically two major pathogens in pigs and they were zoonotic pathogens. There was extensive cellular tropism for both PCV2 and SS2, as well as, PK15 cells was PCV2 infection mainly cell model. It was found that when PCV2 infected PK15 cells before at the MOI=0.1, SS2 could cause more damage to PK15 cells. ITRAQ labeling proteomic technology was used to explore the differentially expressed proteome of PCV2 and SS2 single infection and co-infection in PK15 cells. The results showed that there were total 4736 proteins distinct changed in this infection models for PK15 cells. PCV2 aggravated SS2 infection were showed directly in the big amount of differentially expressed proteins like AGO3, OSBPL1A, ALB, RIC8A, UBL4A, INTS5 and so on. For the KEGG pathway analysis, it indicated that PCV2 mainly induced the proteins lessen though a series of disease pathway like metabolic pathways, huntington’s disease, insulin signaling pathways, long-term depression, etc. to make cell at a state of compensation. PCV2 before infection made the cells was on the chopping block. Contemporary, SS2 induced the proteins of PK15 rapidly changing, including pathways like spliceosome, endoplasmic reticulum, tight junction, actin cytoskeleton and some involved in parasitic infections pathways like ECM-receptor interaction, Leishmaniasis, Toxoplasmosis and so on. Simultaneously, PCV2 and SS2 existed common ground that they influence PK15 cell by some virulence factor interacted with cytomembrane, influenced the function of ribosome and tRNA. The role of SS2 in the co-infection like a cook chopper.

## INTRODUCTION

Porcine circovirus type 2 (PCV2) was a single stranded, non-capsular, closed circular DNA virus including 1,766-1,768 nucleotides ^1–3^. PCV2 was pivotal pathogen, involving in porcine circovirus related disease, porcine dermatitis and nephropathy syndrome, PCV2 reproductive disease, PCV2 lung disease, severe systemic PCV2 infection and PCV2 enteric disease ^4–12^. Porcine circovirus-related diseases were considered to be multifactorially affected diseases in the swine industry ^13^, of which can promote co-infection with other viruses or bacteria ^14–16^, causing aggravation or sharpness of the disease ^4^. PCV2 can damage the pig’s immune system, characterized by lymphadenopathy, lymphopenia, apoptosis and autophagy, leading to severe immunosuppression, causing huge chance to the other agent infection ^17–20^. At the same time, PCV2 could lead to zoonotic disease ^21,22^ and kept persistently infection in PK15 cells ^23^. Streptococcus suis (S. suis) was a Gram-positive facultative anaerobic bacterium, of which type 2 was most serious pathogen of pigs and could be transmitted to humans through close contact with pigs ^24,25^. It was the cause of porcine polyserositis and other diseases, such as heart inflammation, brain infection, bronchial inflammation, arthritis syndrome, sepsis and abortion etc. ^26–30^. According to the clinical detection data, the infection rates of the two pathogens in the healthy and sick pigs were 17% and 35%, respectively, accounting for 69.5% and 59.4% of the total infection or co-infection of the two pathogens, indicating that co-infection of the two pathogens was more common in the clinic in China ^31^. The co-infection of the two pathogens was also reported in Korea, but the symptom was atypical like PCAD ^32^. When pathogens infected host cells, the progression of infection depended on host-pathogen interaction that were regulated in time and space, nevertheless, this response was complex, and the most intuitive performance was the differential proteins changed in host cells ^33,34^. These interactions represent anti-pathogenic and pro-pathogenic cellular responses. So, proteomics study about the host-pathogen involves many aspects. First, we should to explore the differential proteins between treatment and control groups. As the progress of protein labeling and quantification techniques combine with Mass spectrometry (MS), like, label-free MS quantification, which was easy, multifunctional, and could be applied to any biological system. Successively, the stable isotope labeling of amino acids in cell culture (SILAC) ^35^, tandem mass tags (TMT) ^36^ or other isobaric tags trace quantification (iTRAQ) ^36^ were emerge and mature applied in pathogen and host. In studying host-virus interactions, SILAC was employed to study hepatitis C virus ^37^. Proteomics study of porcine circovirus types 2 and 3 on macrophages and porcine lung tissue has been reported ^38,39^. For bacteria, IP-MS was used to identify interactions between effector proteins secreted by intracellular Salmonella and host proteins, and SILAC quantification helped assess specificity of interactions ^40^ and ITRAQ was used to vaccine selection for Mycoplasma bovis ^41^. At present, vaccines were mainly used to prevent and treat viral diseases and antibiotics were used to treat bacterial diseases. However, there was still a lack of drugs for the treatment of viral diseases, meanwhile, the mutation rate of the virus was very quickly, and bacterial resistance was also a serious problem, bacterial disease treatment was still a big challenge. As organism was always exposed to bacteria and virus environment, the changes of the organism should be studied to understand the pathogenic mechanism of viruses and bacteria and to provide theoretical guidance for the prevention and treatment ^42^.

## MATERIALS AND METHODS

### Cells, viruses, bacteria and antibodies

The PCV-free porcine kidney epithelial cell lines (PK15) and PCV2 (10^5.75^ TCID_50_ /ml) strain, which frozen in liquid nitrogen at -80°C, respectively, were kept in Jiangsu Academy of Agricultural Sciences -Institute of Veterinary Medicine (JAAS-IVM) laboratory and maintained in Dulbecco’s Modified Eagle Media (DMEM; Gibco, Carlsbad, CA) containing 7% gamma-irradiated fetal bovine serum (FBS, Gibco). SS2-1 stored in 50% glycerol was kept in JAAS-IVM. Bacterial cultured in Todd Hewitt Broth (THB) medium at 37°C 200 r/min. The polyclonal antibody (pAb) and monoclonal antibody (mAb) against the PCV2 Cap protein and SS2 were obtained as previously and the working concentration was used in 1:1000 and 1:400 dilution, respectively. The polyclonal antibody DExH-box helicase 9 (DHX9) (ABclonal), RIC8A guanine nucleotide exchange factor A (RIC8A) (ABclonal), Annexin 4 (ANXA4) (ABclonal), L-lactate dehydrogenase (LDHA) (ABclonal) were used in 1:1000 dilution, and mAb proliferating cell nuclear antigen (PCNA) (abcam) and glyceraldehyde-3-phosphate dehydrogenase (GAPDH) (ABclonal) were used in 1:10000 dilution.

### Cell culture, virus and bacterial inoculation

There were 8 groups. The 1st and 2nd groups were the blank cells with no deal with for cultured with 50h and 54h named Blank-2h and Blank-6h. The 3rd and 4th groups were the blank cells infected with PCV2 (MOI=0.1) cultured with 50h and 54h named PCV2-2h and PCV2-6h. The 5th and 6th groups were the blank cells cultured with 48h and then infected with SS2 (MOI=10) with 2h and 6h named SS2-2h and SS2-6h. The 7th and 8th groups were the blank cells infected with PCV2 (MOI=0.1) for 48h and then infected with SS2 (MOI=10) for 2h and 6h named PCV2-SS2-2h and PCV2-SS2-6h. PK15 cells were passaged to 3rd generation and maintained at 37°C with 5% CO_2_ for different use. After 48 hours, the culture conditions of the cells were in DMEM with 7% FBS and the virus was in DMEM with 2% FBS.

### The cell viability assay

PK15 cells inoculated with PCV2 (MOI=0.1) was prepared for 48h earlier, the same as blank PK15 cells. PK15 cells was infected with different dose (MOI=0.1, 1, 2.5, 5 and 10) and different co-culture times (2h, 4h, 6h, 12h, and 16h) for SS2. Then CCK8 solution was added 10µL in each well. In order to permitted the formazan dye adhere to the alive cell and reactive, each plate was placed at 37 °C for 2h. For detection, the cell plate was shaken for 10min. Then, a microplate reader (Bio-Rad, model 450) was used to measure its absorbance of each well. Cell viability assay at the formula:

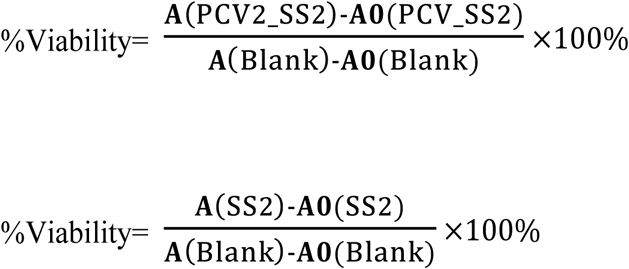

**A**(PCV2_SS2): the absorbance of PK15 cell co-infected with PCV2-SS2

**A0**(PCV2_SS2): the absorbance of the only cell culture fluid co-infected with PCV2-SS2

**A**(SS2): the absorbance of PK15 cell co-infected with SS2

**A0**(PCV2_SS2): the absorbance of the only cell culture fluid co-infected with SS2

**A**(Blank): the absorbance of PK15 cell without pathogen infection

**A0**(Blank): the absorbance of the only cell culture fluid

### Determination of iTRAQ experimental protocol

The different group samples were prepared according to experimental design mentioned in preceding part of this text. In order to achieve the comparison and inspections of different groups. The overall design of proteomics was shown in Figure 2. A certain quantity sample was taken from each group of the 8 groups in a ratio of 1:1…1:1 for mixing and forming a new group. The 9 groups were marked separately and reclassified into two new groups for iTRAQ assay according to the time 2 hours and 6 hours. This was done to make it easier to compare changes between groups.

### Protein preparation

Different group samples were added with protein lysate (7M Urea, 2M Thiourea, 4% SDS, 40mM Tris-HCl pH8.5, 1mM PMSF, 2mM EDTA) mixing and 5 minutes in ice, then, added a final concentration of 10mM DTT in ultrasound and ice for 15min. The treated samples were centrifuged at 13000g, 4°C for 20 minutes. The different supernatants were picked in new centrifuge tubes, added cold acetone overnight at -20°C, then centrifuged at 13000g, 4°C for 20 minutes and collecting sediment and dried in air. The treated samples dissolved in 8M Urea, 100mM TEAB (pH8.0) and added DTT to a final concentration of 10mM, water bath in 56°C for 30 minutes. Subsequently, the samples added IAM to a final concentration of 55 mM putting in dark place at room temperature for 30 min. Bradford method was used to measure of protein concentration.

### Digestion and desalting

The same amount of protein from each sample was used for trypsin digestion. For each sample, 100µg of protein was diluted 5 times, and then added trypsin at a mass ratio of 1:50 (trypsin: protein), digesting overnight at 37°C. The samples after enzymatic hydrolysis was desalted on a C18 column, and the dehydrated peptide was freeze-dried under vacuum.

### iTRAQ labeling and fractionation

The dried peptide was solubilized with 0.5M 20uL TEAB for peptides labeling. Samples were labeled with iTRAQ Reagent-8 plex Multiplex Kit (AB Sciex U.K. Limited) according to the manufacturer instructions. All of the labeled samples were mixed with equal amount. Then, the labeled samples were fractionated using high performance liquid chromatography (HPLC) system (Thermo DINOEX Ultimate 3000 BioRS) using a Durashell C18(5 um, 100Å, 4.6 × 250 mm). Peptides separation was achieved using a gradually increasing ACN concentration under alkaline conditions at a flow rate of 1 ml/min and one tube was collected per minute. A total of 42 secondary fractions were collected and combined into 12 fractions. The combined fractions were desalted on a Strata-X column and dried under vacuum.

### Proteomics LC-MS/MS Analysis

LC-ESI-MS/MS analysis was performed on an AB SCIEX nanoLC-MS/MS (Triple TOF 5600 plus) system. Samples were chromatographed using a 90min gradient from 2–98% (buffer A 2% (v/v) acetonitrile and 0.1% (v/v) formic acid, buffer B 98% (v/v) acetonitrile, and 0.1% (v/v) formic acid) after direct injection onto a 20µm PicoFrit emitter (New Objective) packed to 12cm with Strata-X C18 AQ 3µm 120 Å stationary phase. For IDA (Information Dependent Acquisition), MSI was performed with fragmentation accumulation time of 250ms, and MS2 collected 30 precursor ions with fragmention accumulation time of 50ms. MS1 and MS2 collected the range 350-1500m/z and the range of 100-1500m/z, respectively. The precursors fragmentation was excluded from reselection for the first 15s.

### Proteomics Data Analysis

ProteinPilot^TM^ Software V4.5 was used to acquisition the original data file from MS/MS directly. The Paragon algorithm2 was integrated into ProteinPilot and employed against Uniprot Sus scrofa species (PR1-18110005 items, update in Nov. 2018) for database searching and for protein identification. The parameters were set as follows: the instrument was TripleTOF 5600, iTRAQ quantification, cysteine modified with iodoacetamide; biological modifications were selected as ID focus, trypsin digestion, the Quantitate, Bias Correction and Background Correction was checked for protein quantification and normalization. Proteinpilot has taken into account all possible modifications in the search, while adding automatic fault decoy matching function, using the PSPEP (Proteomics System Performance Evaluation Pipeline Software, integrated in the ProteinPilot Software) algorithm. Only proteins with at least one unique peptide and unused value more than 1.3 (the reliability level is above 95%) were identified. These identified peptides were excluded from the quantitative analysis for: (i) peptides missing an iTRAQ reagent label, (ii) peptides peaks did not match the corresponding iTRAQ labels, (iii) peptides with too low S/N (signal-to-noise ratio), (iv) peptides with low identification confidence, (v) peptides were not unique peptide. Gene Ontology (GO, http://geneontology.org/) is an internationally standardized gene function classification system that provides a dynamically updated of new vocabulary to fully describe the properties of genes and gene products in an organism. There was a total of three ontology that describe the molecular function, cellular component, and biological process of the gene in GO, respectively. When protein abundance ratio, measured in iTRAQ after normalized (P-value≤0.05 for corresponding screening), the protein was considered to be a significantly changed protein between different groups. The data of identified proteins was then selected by the criterion (up-regulate≥1.5 or down-regulate≤0.67, P-value≤0.05) for next analysis. The analysis mapped the changed proteins to the various terms from the Gene Ontology database and calculates the number of proteins per term, then, functional annotation for significant changed proteins of GO term. Cluster of Orthologous Groups of proteins (COG, http://www.ncbi.nlm.nih.gov/COG/) is a database of orthologous classification of proteins. The proteins of COG were assumed to be derived from an ancestral protein, therefore, either orthologs or paralogs. Orthologs refers to proteins that have evolved from vertical pedigrees from different species, and retain the same typically function of the original protein. Paralogs are proteins that are derived from gene duplication of a certain species and may evolve into a new and the same as the previously relevant functions protein. To identify candidate biomarkers, we employed hypergeometric test to perform GO enrichment and KEGG pathway enrichment. To point out the protein-protein interactions, Search Tool for the Retrieval of Interacting Genes/Proteins (STRING) was employed (http://www.string-db.org/). All other pictures were drawn with R language (http://www.r-project.org/).

### Venn diagram

Venn analysis was carried out for different comparison groups of the identified significant differential proteins and differential proteins involved in signaling pathways. Venn diagrams were constructed online (http://bioinfogp.cnb.csic.es/tools/venny/index.html) according to the introduction on the website.

### Ingenuity Pathways Analysis (IPA)

To obtain further insight into the functions of the identified distinct abundant proteins, pathways and molecular networks between groups were analyzed using IPA (Ingenuity Systems, Redwood City, CA). Briefly, identified proteins were uploaded to the IPA software and mapped to their corresponding gene objects, and biological functions, canonical pathways and molecular networks of gene objects were subsequently evaluated and constructed based on information in the IPA database. Statistics for the functional analysis were calculated automatically by the software using the right-tailed Fisher’s exact test. Molecular networks generated based on ranking scores were optimized to include as many proteins from input abundance profiles as possible, and to maximize network connections. Nodes in networks were coloured, and different shapes were used to represent fold changes and functional classes of genes and gene products. Interactions between nodes were connected only when supported by at least one or more references.

### Relative quantitative PCR (qPCR)

The proteins (changed in abundance and had a significant impact on co-infection) identified by iTRAQ were picked up and found the corresponding gene conserved sequences corresponding to a gene. To validate the changes in abundance of the proteins identified by iTRAQ, total mRNA was extracted from cells infected with one or both viruses using TRIzol reagent (Magen) and reverse-transcribed using the SuperScript First-Strand Synthesis System (Fermentas, Pittsburgh, PA) according to manufacturer instructions. The GAPDH gene was used as an internal standard, and the relative quantitative abundance of selected genes was assayed by real-time PCR using the SYBR Premix Ex Taq (ABclonal, WuHan, China) with an ABI 7500 sequence detection system (Applied Biosystems). The results were calculated as the mean value of triplicate reactions. Primers used for amplification of the selected genes were shown in Table S-1.

### Western Blot Assay

Cells were washed with PBS two times. Then treated with RIPA (EpiZyme) according to the protocol (http://www.epizyme.cn/index.php?c=article&id=702). Cells were lysed in ice for 30 minutes and collected the supernatant. The protein sample was quantified with the BCA kit and the mass action concentration was 3.5ug/uL. The samples were mixed with 5 plus protein loading buffer and heated in 100°C for 5 minutes and separated on a 12% SDS-PAGE (10µL per lane). After electrophoresis, proteins were transfered to a PVDF membrane (Millipore) at 23 volts for 0.5 hour. The PVDF membranes were blocked in PBS with 5% skim milk for 1 hour. The membranes were incubated with the ANXA4, LDHA, DHX9, RIC8A (ABclonal) rabbit anti-human antibody (with the dilution of 1:1000) overnight at 4 °C and PCNA (rabbit anti-human) (abcam) and GAPDH mouse anti-human (ABclonal) and then with the secondary antibody HRP labeled goat anti-rabbit IgG (with the dilution of 1:5000, ABclonal) for 1 hour. Protein levels were monitored by enhanced chemiluminescence. PNCA and GADPH were employed as protein loading control.

### Immunofluorescent Staining

The 8 treatment groups in PK15 cells were stained with the following primary antibodies: rabbit anti-PCV2 (Abcam), rabbit anti-SS2-1. Treatment groups were then incubated with secondary antibodies with FITC and CY3. All images were captured on a Nikon fluorescent microscope equipped with a digital camera (TE2000-U, Nikon). Each immunofluorescent stain was repeated three times using serial sections; negative controls were included to determine the amount of background staining.

### Statistical method

Data were presented as mean ± SD of at least three individual experiments. Statistical differences between groups were analyzed by the unpaired t-test using Microsoft Excel 2013. A value of P <0.05 was considered statistical significance. Graphpad prism 7 was used to processing composite image and drawing figures.

### Ethics statement

The procedures used for the preparation of pAb and mAb in experimental animals were approved by the Institutional Animal Care and Use Committee (IACUC) of Jiangsu Academy of Agricultural Sciences (Permit No. SYXK 2015-0020), in accordance with the Regulations for the Administration of Affairs Concerning Experimental Animals approved by the State Council of PR China.

## RESULTS

### Cell viability assay

PK15 cells inoculated with PCV2 (MOI=0.1) was prepared for 48h earlier, the same as blank PK15 cells. PK15 cells was infected with different dose (MOI=0.1, 1, 2.5, 5 and 10) and different co-culture times (2h, 4h, 6h, 12h, and 16h) of SS2 (Figure 1). The results showed that the survival rate of PK15 cells was not exceed 81.9% and 78.4% within 2 hours, and 66.2% and 57.6% within 6 hours of SS2 infection and PCV2-SS2 co-infection at the MOI=10. This figure also showed that the damage of SS2 was dose dependent and time dependent. The cell basically died after 16 hours. The most distinctive feature of this results indicated that PCV2 infection promoted the damage of SS2 to PK15 cells at the corresponding time and MOI. Basic to the result, PK15 cells, (infected with PCV2 at the MOI=1 with 48 hours and SS2-1 at the MOI=10 with 2 and 6 hours) were selected to further proteomics study.

**Figure 1.**
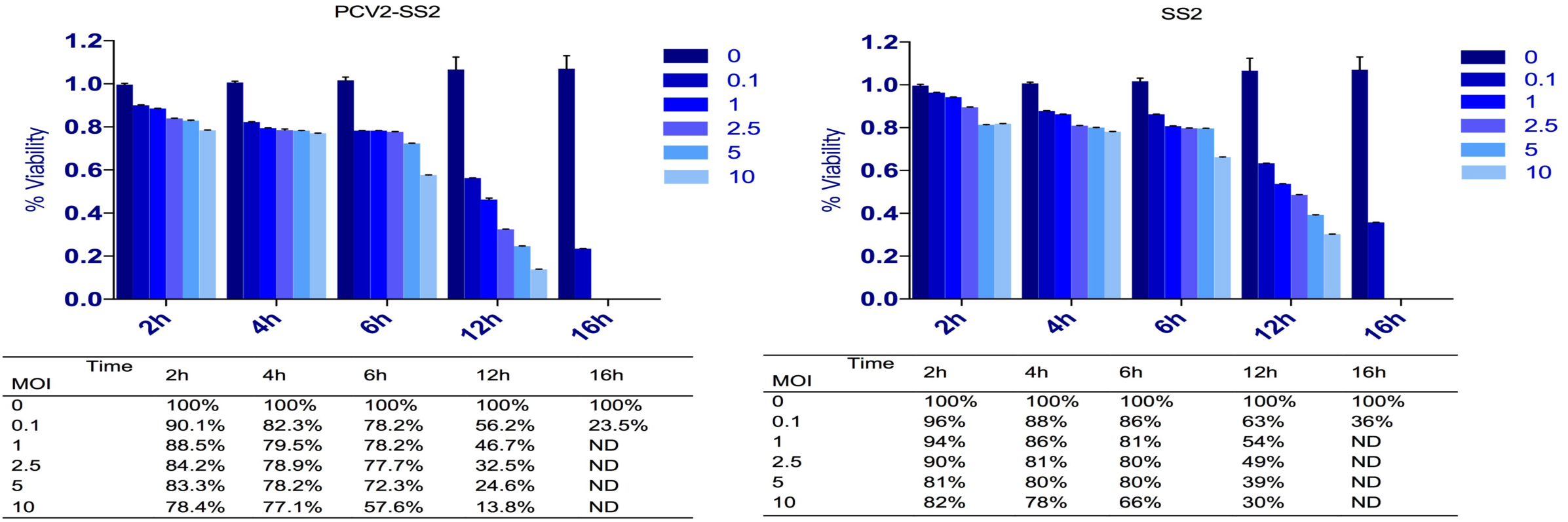
The cell viability assay when SS2 single infection and with PCV2 co-infection. The ordinate indicated the cell survival rate/viability, and the abscissa was the time that SS2 interaction with PK15. The infection MOI of SS2 was divided into six groups, they were 0, 0.1, 1, 2.5, 5, and 10, respectively, represented by dark to light blue.

**Figure 2.**
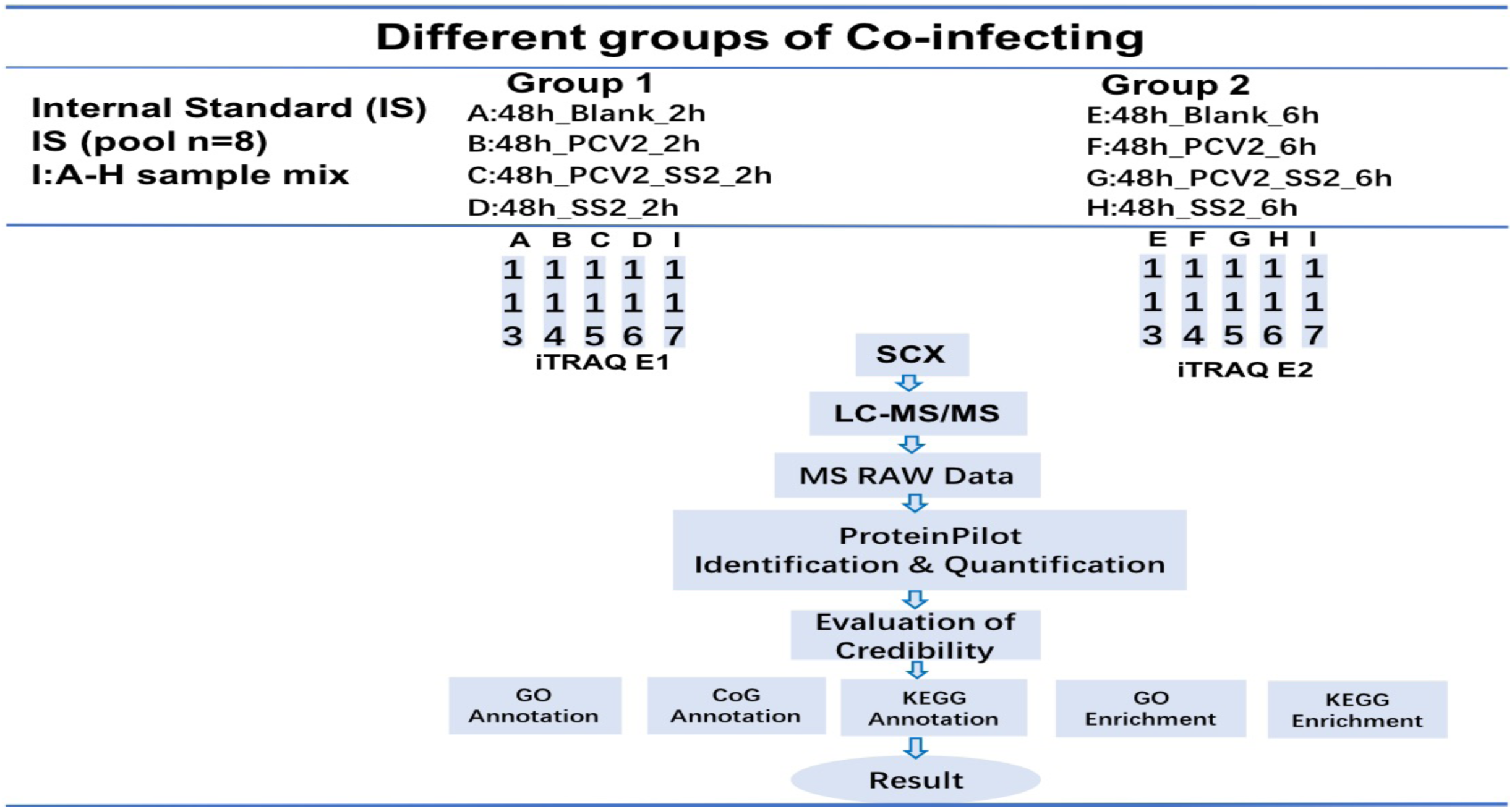
Proteomics implementation schemes for different groups using iTRAQ. There were A, B, C, D, E, F, G and H 8 groups were ready to be used for iTRAQ and proteomics analysis. The same amounts of samples were extracted from these 8 groups to form an internal standard group named I. These 9 groups conducted two ITRAQ experiments named ITRAQ E1 and E2 which all including I group. And then the experiments were analyzed in accordance with the procedure.

### Identification and quantification of distinct abundant proteins by using iTRAQ-based LC-MS/MS

The MS/MS spectra of unique peptides infer to the identified putative distinct abundant proteins PCNA, GAPDH, ANXA4, DHX9, LDHA, RIC8A were shown in (Figure S-1). A total of 4,736 proteins were identified in this project. Due to the limitations of the background date annotation library, it was not all proteins in Uniprot database has annotation information. There were 4,602 proteins in GO function annotation, 2457 proteins in COG function annotation and 2569 proteins in KEGG function annotation (Figure 3 (a)). This experiment provided GO annotations, KEGG annotations and COG annotations to fully reflect the function of proteins using different sources of databases, revealing the biological significance of proteins in various life activities (Tables S-2 to S-4, respectively).

**Figure 3.**
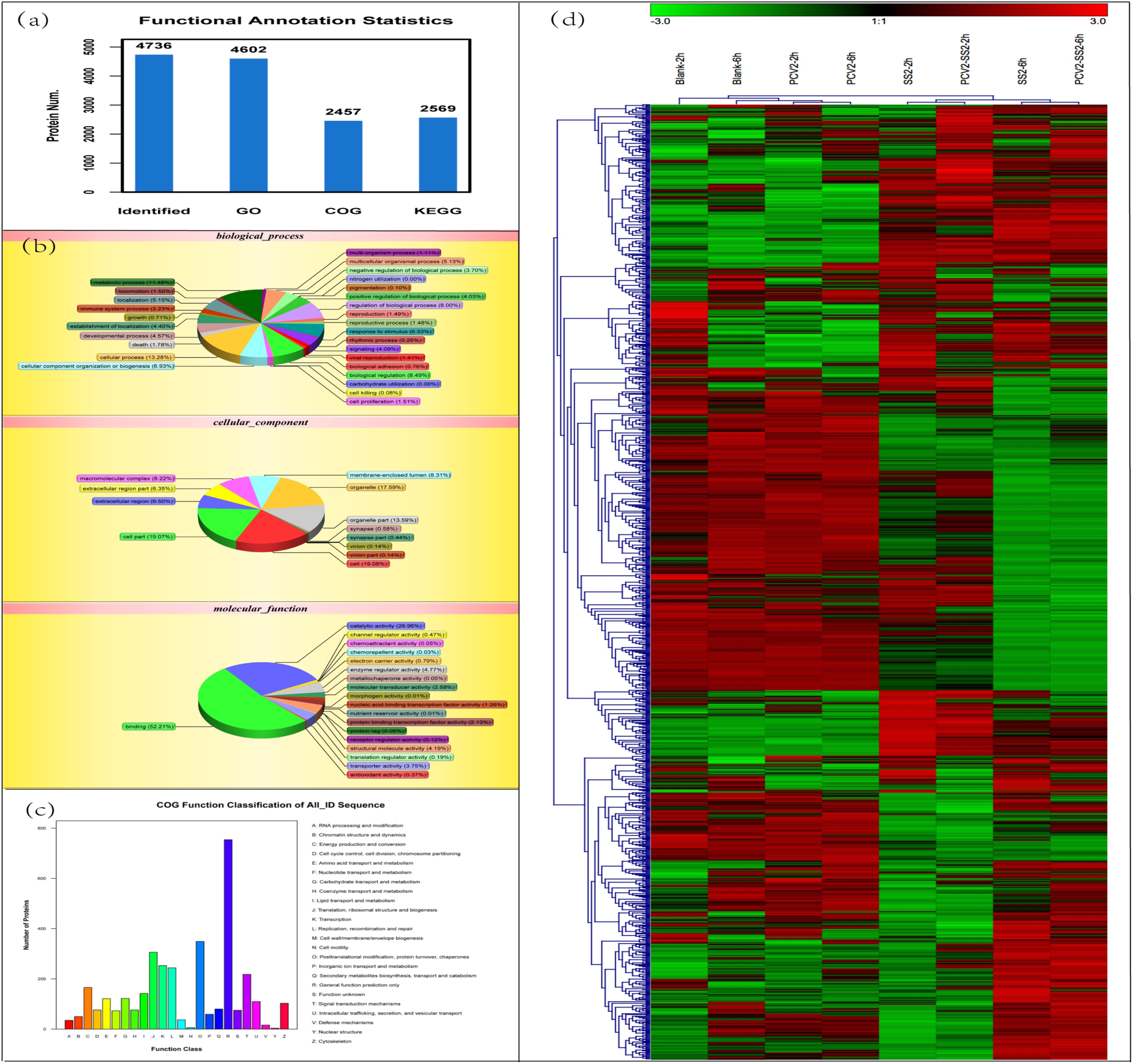
Number of total proteins in GO, COG and KEGG database, distribution of total proteins in GO and COG and cluster analysis of total proteins for 8 groups. (a), the total proteins identified in GO, COG, and KEGG database. (b), GO analysis of 4602 proteins including biological process, cellular process and molecular function which also including different secondary functional classifications and proportions marked in different colors. (c), COG function classification of 2457 proteins and different colors represents different COG functions and proportions. (d), the cluster analysis of identified proteins in 8 groups, of which the ordinate indicated the near and far relationships according unique peptide of different identified proteins and the abscissa indicated proximity of different groups. Green color represented proteins down-regulated, red color represented proteins up-regulated and the color shade degree indicated the degree of up and down regulated.

### The expression level hierarchical clustering analysis

Hierarchical clustering analysis brought the higher similarity proteins together intuitively. Clustering the samples together by column, it could not only reflect the repeatability of experimental design, but also judge the correlation between samples of different groups. This analysis could provide the information that in a special condition, one or same samples with high content or low content or even 0 of one certain or some proteins. The results showed all the 1184 significant difference proteins of compilations of the 8 different groups (samples) (Figure 3 (d)). This horizontal of hierarchical clustering analysis showed that, according to the degree of protein proximity, the order of different groups was Blank-2h, Blank-6h, PCV2-2h, PCV2-6h, SS2-2h, PCV2-SS2-2h, SS2 and PCV2-SS2-6h. Simultaneously, it reflected the similarity degree of proteins between different treatment groups. Among them, 2 blank groups were closed, 2 PCV2 single infected groups (with the treatment time 50 hours and 54 hours) were closed and the SS2, PCV2-SS2 treatment groups (with the treatment time 2 hours and 6 hours) were closed. The vertical 1184 proteins were hierarchical clustered according to the similarity correlations and the proteins ID (UniProt) were listed in Figure S-2. Proteins in the same cluster showed similar abundance, similar characteristics, or participate in the same cellular process.

### GO annotation

We performed a GO functional annotation analysis on all identified proteins, and for the GO class involved in the three ontologies (cellular component, biological process, molecular function). At the same time, make a chart (Figure 3 (b)) and omit the GO class without the corresponding protein. For different groups/samples Blank-2h, Blank-6h, PCV2-2h, PCV2-6h, SS2-2h, SS2-6h, PCV2-SS2-2h and PCV2-SS2-6h, GO annotation were performed GO annotation analysis to compare the treatment groups on differential proteins like PCV2-2h: Blank-2h, PCV2-SS2-2h: Blank-2h, SS2-2h: Blank-2h, PCV2-6h: Blank-6h, PCV2-SS2-6h: Blank-6h and SS2-6h: Blank-6h (Figure 4). As showed in Figure 4, as stated in GO classification, all identified proteins in different groups were generally divided into three classes (biological process, cellular component, and molecular function) and further subcategorized into 45 hierarchically structured GO classifications for detailed investigation. It showed the proportion of up-regulated and down-regulated proteins in each subcategory. The results showed that biological process contained cellular process, metabolic process, biological regulation, regulation of biological process, cellular component organization or biogenesis, response to stimulus, multicellular organism process, development process, localization, negative regulation of biological process, establishment of localization, positive regulation of biological process, signaling, immune system process, death, locomotion, reproduction, reproductive process, cell proliferation, biological adhesion, growth, viral reproduction, multi-organism process, cell killing, rhythmic process, while cellular contained cell, cell part, organelle, macromolecular complex, membrane enclosed lumen, extracellular region, extracellular region part, synapse, synapse part and molecular function contained binding, catalytic activity, enzyme regulator activity, transporter activity, structure molecular activity, molecular transducer activity, antioxidant activity, channel regulator activity, electron carrier activity, translation regulator activity. For different groups, the distribution of major components was similarity, but not the same.

**Figure 4.**
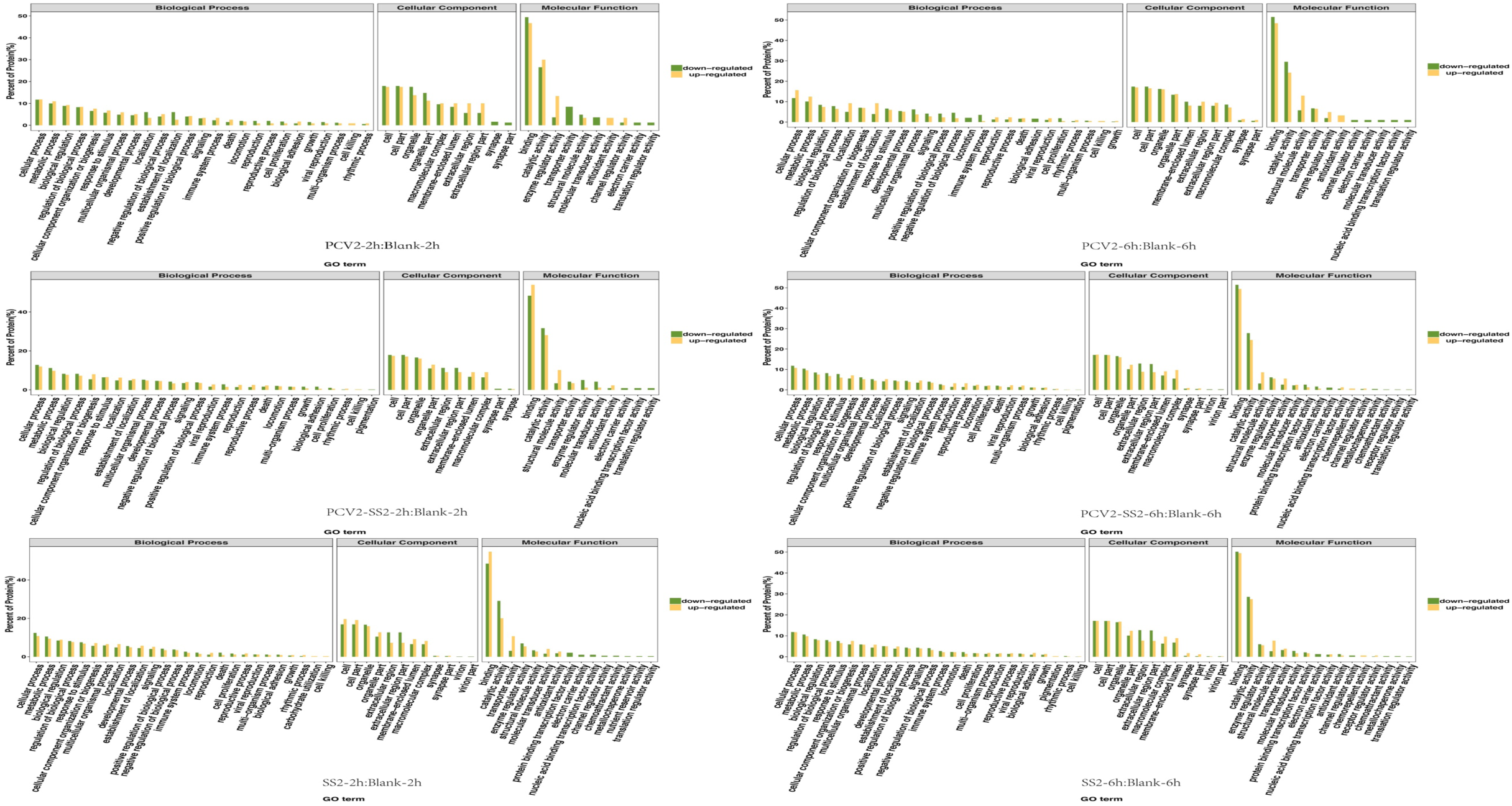
GO analysis of differential proteins in treatment groups to blank groups. The ordinate indicated the percentage of protein, the functional classifications in biological process, cellular component and molecular function, respectively. Green color and yellow color represented down and up regulated, respectively. The groups included PCV2-2h:Blank-2h, PCV2-6hBlank-6h, PCV2-SS2-2h:Blank-2h, PCV2-SS2-6h:Blank-6h, SS2-2h:Blank-2h and SS2-6h:Blank-6h.

### COG annotation

We compared the identified proteins with the COG database, predict the possible functions of these proteins and perform functional classification statistics on them. Figure 3 (c) indicated all these proteins showed 24 functions, and the numbers is Tables S-4. The most abundant proteins were posttranslational modification, protein turnover, chaperones proteins and the least abundant proteins were nuclear structure proteins. This result provoked us to focus on the organelle and organelle part relevant differential expressed proteins.

### GO pathway analysis

All the identified proteins had been performed pathway analysis according to the KEGG database and found the most import ant biochemical metabolic pathways and signal transduction pathways involved in protein. Figure 5 shows the metabolic pathways and signal transduction pathways of proteins between different groups. It also illustrated the ratio of up-regulated and down-regulated proteins to different metabolic and signal transduction pathways. This result showed that metabolic pathways is the mainly pathways among these groups. In PCV2 groups, the mainly up-regulated and down-regulated pathways existed big difference. The up-regulated proteins involved in metabolic pathways, glycolysis/gluconeogenesis, microbial metabolism in diverse environments, aminoacyl-tRNA biosynthesis, oocyte meiosis, pentose phosphate pathway, proteasome, Wnt signaling pathway, long-term depression, aldosterone regulated sodium reabsorption, ribosome, purine metabolism, pyruvate metabolism, cysteine and methionine metabolism, and pyrimidine metabolism, while the down-regulated proteins involved in metabolic pathways, huntington’s disease, lysine degradation, insulin signaling pathway, phagosome, microbial metabolism in diverse environments, endocytosis, oxidative phosphorylation, spliceosome, gastric acid secretion, protein processing in endoplasmic reticulum, focal adhesion, ECM-receptor interaction, pathway in cancer, Alzheimer’s disease, toxoplasmosis, small cell lung cancer and cell adhesion molecules. In PCV2-SS2 groups, the up-regulated proteins involved in metabolic pathways, ribosome, regulation of actin cytoskeleton, tight junction, microbial metabolism in diverse environments, protein processing in endoplasmic reticulum, spliceosome, alanine, aspartate and glutamate metabolism peroxisome, pathogenic Escherichia coli infection, aminoacyl-tRNA biosynthesis, pathways in cancer, toxoplasmosis, protein processing in endoplasmic reticulum, complement and coagulation cascades, focal adhesion and amoebiasis, while the down-regulated proteins involved in metabolic pathways, microbial metabolism in diverse environments, focal adhesion, regulation of actin cytoskeleton, ECM-receptor interaction, RNA transport, spliceosome, dilated cardiomyopathy, arrhythmogenic right ventricular cardiomyopathy (ARVC), hypertrophic cardiomyopathy, glycolysis/gluconeogenesis, regulation of actin cytoskeleton, purine metabolism, pyruvate metabolism, lysosome, pentose phosphate pathway, cysteine and methionine metabolism. In SS2 groups, the up-regulated proteins involved in spliceosome, protein processing in endoplasmic reticulum, metabolic pathways, staphylococcus aureus infection, ubiquitin mediated proteolysis, complement and coagulation cascades, aminoacyl-tRNA biosynthesis, malaria, tight junction and regulation of actin cytoskeleton and pathway in cancer, ECM-recepror interaction, focal adhesion, cell cycle, DNA replication, small cell lung cancer, ramoebiasis, viral myocarditis, while the down-regulated proteins involved in metabolic pathways microbial metabolism in diverse environments, glycolysis/gluconeogenesis, pathogenic Escherichia coli infection, protein processing in endoplasmic reticulum, regulation of actin cytoskeleton, glutathione metabolism, pathways in cancer, endocytosis, tight junction, proteasome, purine metabolism, glutathione metabolism, pyruvate metabolism, cysteine and methionine metabolism. This result showed that there were the same and the different pathways among these main signaling pathways of different groups. The detail information of the differential expression proteins in the major signaling pathways were listed in the attached table S-5. These protein changes objectively and positively reflected the material basis of PCV2 exacerbating the damage of Streptococcus suis to PK15 cells. But how this effect was linked needs further analysis and verification

**Figure 5.**
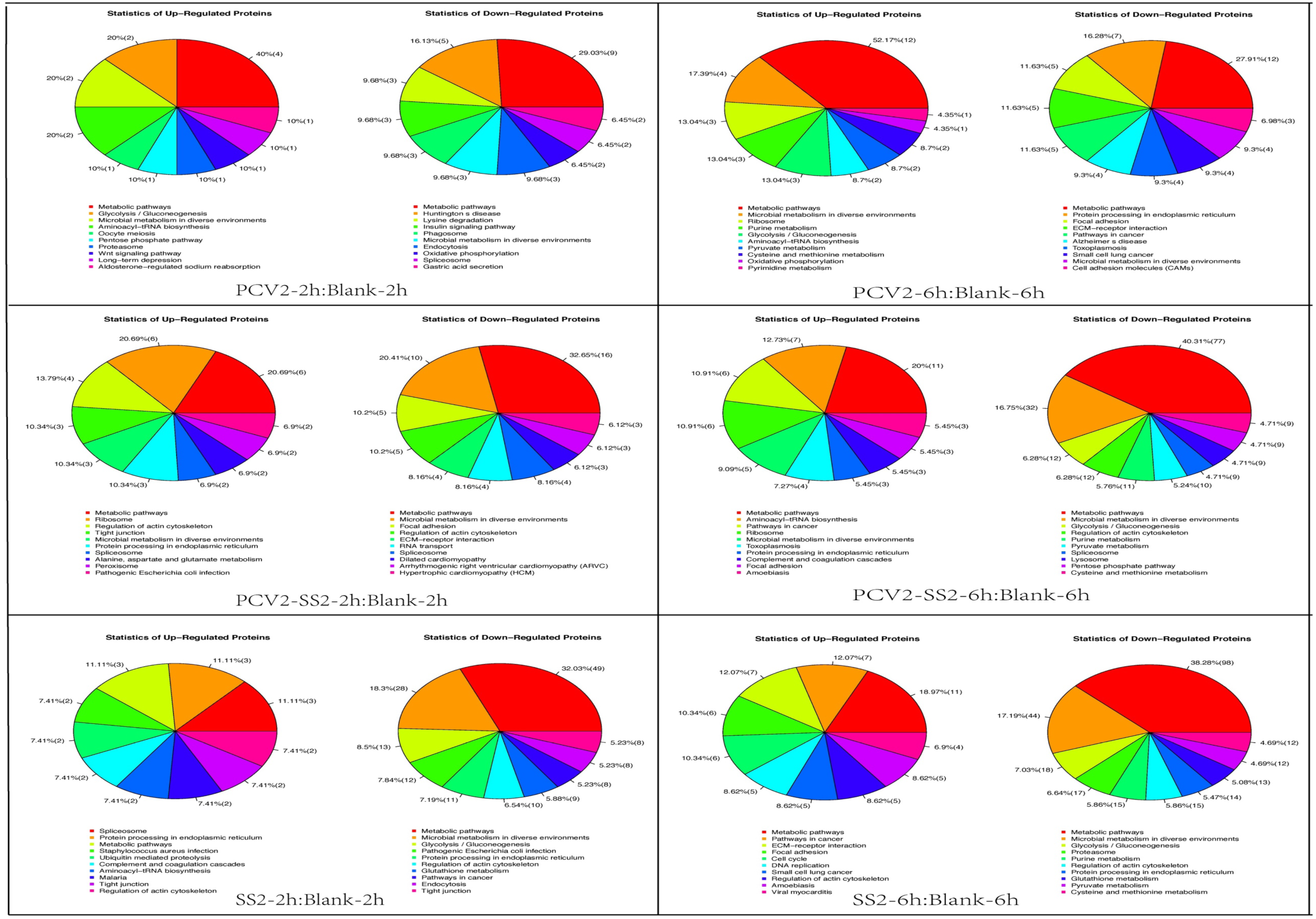
KEGG analysis of differential proteins in treatment groups to blank groups. There were 6 differential proteins groups including PCV2-2h:Blank-2h, PCV2-6hBlank-6h, PCV2-SS2-2h:Blank-2h, PCV2-SS2-6h:Blank-6h, SS2-2h:Blank-2h and SS2-6h:Blank-6h. There were up-regulated and down-regulated differential proteins in the 6 groups. Each pie chart contained the percentage of the top ten major pathways (listed in order from high to low below).

### Venn diagram analysis

In order to further find the connection between these groups, Venn diagram analysis was performed. In other words, identified and pathways involved with significant differences in p2/b2, s2/b2, ps2/b2, p6/b6, s6/b6, and ps6/b6 proteins were analyzed using a Venn diagram (Figure 6). There were 5 common proteins in the treatment of 2 hours groups (p2/b2, s2/b2 and ps2/b2), and they were 40S ribosomal protein S9 (RPS9), Serum albumin (ALB), RIC8 guanine nucleotide exchange factor A (RIC8A), Argonaute 3 (AGO3) and Oxysterol-binding protein (OSBPL1A), which participate in Aminoacyl-tRNA biosynthesis, Ribosome pathway. This result showed that the PCV2 and SS2 could interfere protein synthesis together in the SS2 early infection stage. And there were 15 common proteins in the treatment of 6 hours groups (p6/b6, s6/b6, and ps6/b6), and they were Laminin subunit alpha 5 (LAMA5), Uncharacterized protein LAMC1, Laminin subunit beta 1 (LAMB1), L-lactate dehydrogenase (LDHA), Splicing factor proline and glutamine rich (SPFQ), Annexin (ANXA4), Cathepsin D protein Fragment, Carbonyl reductase 3 (CBR3), 60S ribosomal protein L6 (RPL6), Integrin beta (ITGB1), Serum albumin (ALB), Aminoacylase-1 (ACY1), Cathepsin D protein, Gam ma-tubulin complex component (TUBGCP2), Replication initiator 1 (REPIN1), Golgin A3 (GOLGA3) and Alpha-1,3-mannosyl-glycoprotein 4-beta-N-acetylglucosaminyltransferase B (MGAT4B). This result showed that the 2 pathogens induced more serious cell damage proteins including in Toxoplasmosis, Amoebiasis, Shigellosis, Leishmaniasis, Prion diseases, Pathogenic Escherichia coli infection and Small cell lung cancer pathways after 6 hours infection. Simultaneously, a total of 199 pathways were categorized (Table S-6), among then, 44 same pathways were involved in all 6 groups, 55 same pathways in 5 groups, 36 same pathways in 4 groups, 26 same pathways in 3 groups and 20 same pathways in 2 groups and 20 pathways just in single groups. This showed that the distribution of proteins was very widely in both single infection groups and co-infection groups. And the data indicated that the number of down-regulated proteins was significantly higher than the number of up-regulated proteins in these six treatment groups (Figure 6). This data also could see that the PK15 cells infected with PCV2 before were not as violent reaction as they single infected with SS2, but PCV2 pre-infected cells had lower survival numbers when SS2 invaded.

**Figure 6.**
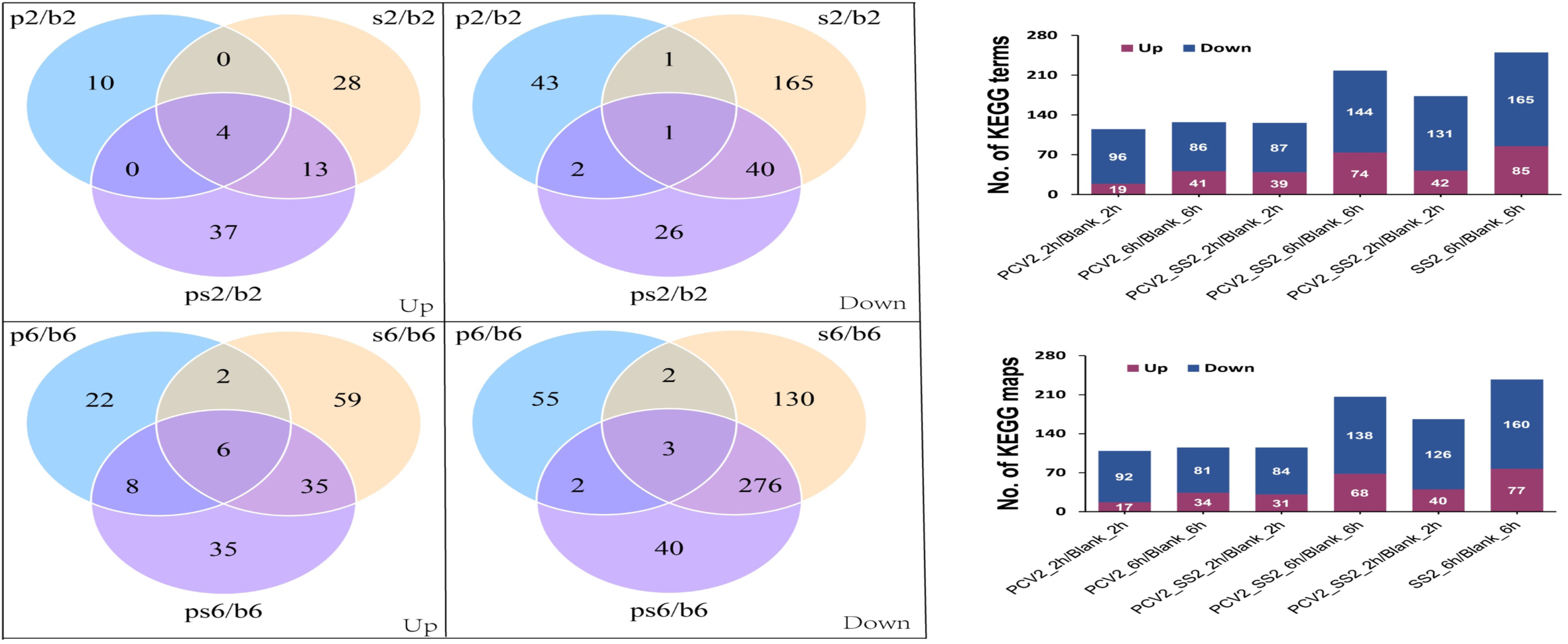
The Venn analysis of differential proteins in treatment groups to blank groups of KEGG data. According to 2 time points of 2h and 6h, the 6 groups of differential proteins were divided into two groups and the up-regulated and down-regulated differential proteins were performed Venn analysis, respectively. The abscissa indicated 6 comparison groups and the ordinate represented the number of proteins involved in pathways and the number of proteins involved in pathways maps, respectively.

### Ingenuity Pathways Analysis (IPA)

In response to the above results, we selected the proteins, related to PCV2 infection and SS2 infection early and intermediate stages, and the preformed STRING analysis to explore protein interaction network (Figure 7) and ingenuity pathways analysis. But some proteins without information in database were not figured. In figure 10, We draw a network diagram that interacts with 40S ribosomal protein S9 (RPS9), RIC8 guanine nucleotide exchange factor A (RIC8A), 60S ribosomal protein L6 (RPL6), Alpha-1,3-mannosyl-glycoprotein 4-beta-N-acetylglucosaminyltransferase B (MGAT4B) and Gamma-tubulin complex component (TUBGCP2). This figure expressed that the interaction relationship in inner cells and force magnitude of interaction of the differential proteins. The proteins, participated in the regulation of PK15 infected by the two pathogens, were draw in corresponding signaling pathway with their up-regulated and down regulated information at the link of (file:///Users/tanjimin/Desktop/pcv2-ss2-pk-ITRAQ%20analysis%20Project_data/2.Enrichment/Pathway_enrichment/PCV2_2h-Blank_2h_Pathway_enrichment/PCV2_2h-Blank_2h.htm) and (file:///Users/tanjimin/Desktop/pcv2-ss2-pk-ITRAQ%20analysis%20Project_data/2.Enrichment/Pathway_enrichment/PCV2_6h-Blank_6h_Pathway_enrichment/PCV2_6h-Blank_6h.htm#gene21).

**Figure 7.**
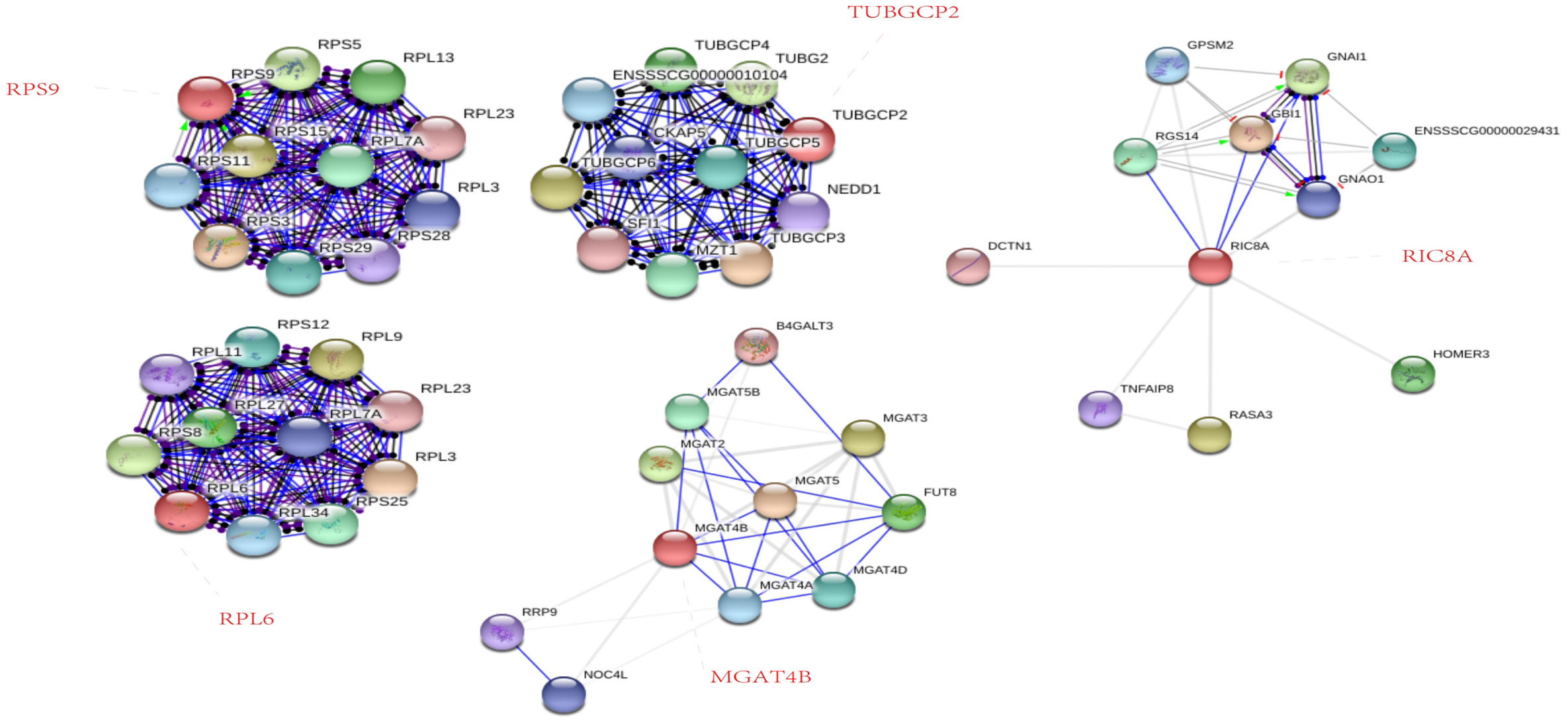
The proteins interaction analysis by STING and the proteins were produced by PCV2 and SS2 co-infection and stimulating PK-15 cells. There were 5 proteins among 14 co-infection proteins existing proteins interaction maps and they were RPS9, TUBGCP2, RPL6, MGAT4B and RIC8A. The force between proteins was represented by different line segments. Network nodes represented proteins. Red nodes represented query proteins and first shell of interactors and white nodes represented second shell of interactors. Empty nodes represented proteins of unknown 3D structure and filled nodes represented some 3D structure was known or predicted. And the edges represented protein-protein associations, the fluorescent green 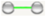 represented activation, the blue 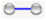 represented binding, the light green 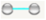 represented phenotype, the dark 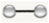 represented reaction, the red 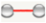 represented inhibition, the purple 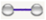 represented catalysis, the light red 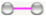 represented posttranslational modification and the yellow 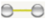 represented transcriptional regulation. The 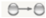 represented positive, the 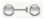 represented negative and the 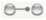 represented unspecified.

This result indicated that PCV2 and SS2 co-infection and the two pathogens single infection to PK15 cells, respectively, there were some key function/pathways participate in. And the common pathways were ribosome and aminoacyl-tRNA biosynthesis. Simultaneously, they also via ECM-receptor interaction pathway interacted with the cell membrane, so as to achieved the purpose of pathogenic. The pathway maps could help to further explore the pathogenic mechanism of the two pathogens.

### Relative quantitative PCR (qPCR)

Based on proteomics data and the comparative analysis. The differential proteins were selected and according to those protein sequences and gene sequences, we designed the relative fluorescence quantitative primers of these genes (Table S-7). PCR results showed that the expression of mRNA was not all the same with the result of proteomics data. The proteins with large variation folds and the reference genes showed consistency with mRNA results trend among different groups. And the iTRAQ data showed distinct differencial proteins were consistency with the qPCR data. This conclusion could be reflected in the results of LDHA, ANXA4, RIC8A, DHX9 mRNA experiments (Figure 8).

**Figure 8.**
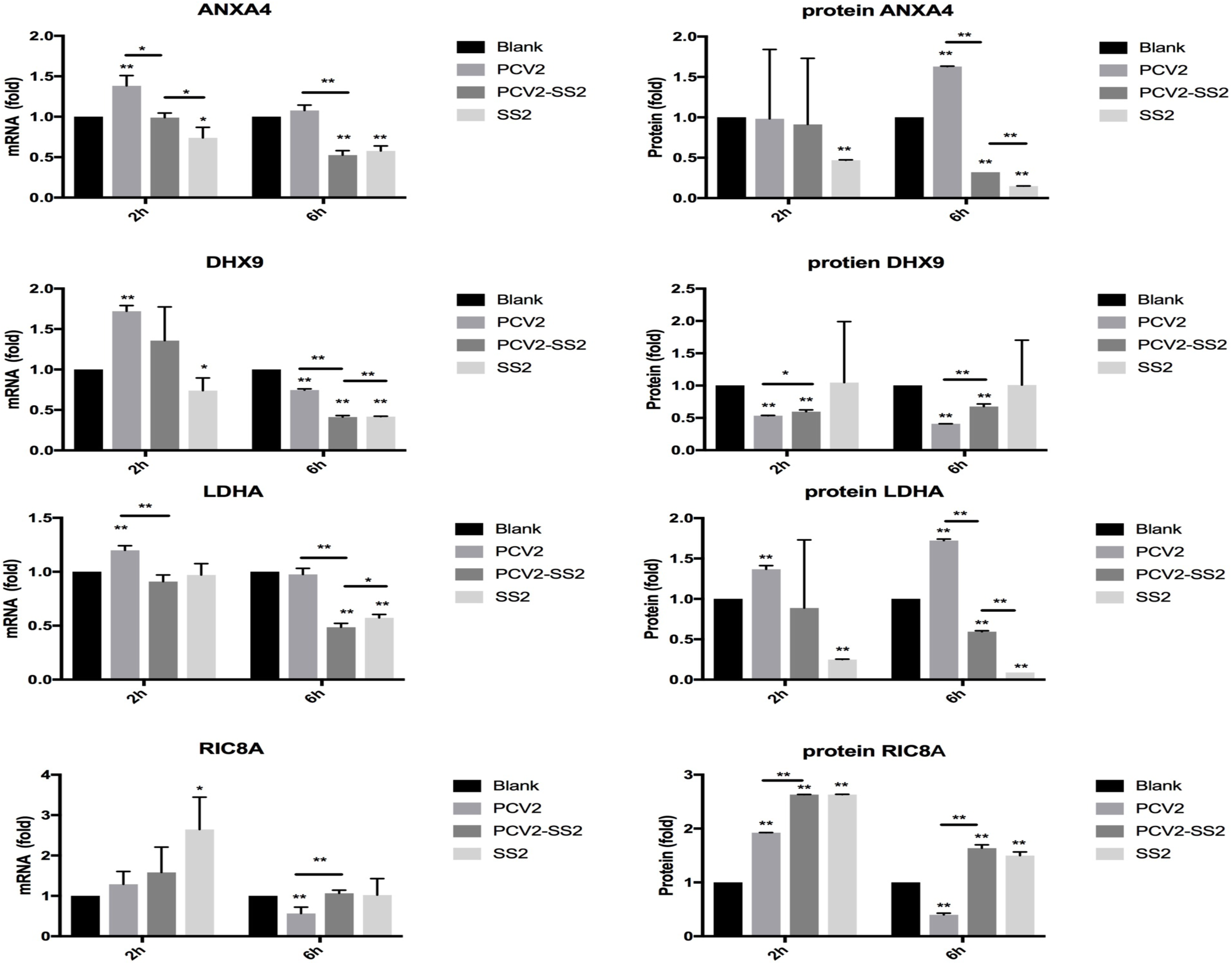
The qPCR verification and the proteins changed folds of iTRAQ data. The left of the figure was qPCR data and the right of the figure was iTRAQ data. The * and **represented p<0.05 and p<0.01. Dark to light gray represented groups Blank, PCV2, PCV2-SS2 and SS2.

### Western Blot Assay

In one aspect, six proteins were detected by western blot to verify the proteomic and bioinformational data (Figure 9). And the results instrumented that the western blot results of LDHA, ANXA4, RIC8A, DHX9 was consistency with the proteomic data and bioinformational data. Simultaneously, this result pointed out several co-infection key proteins of PCV2 infection, SS2 infection, and PCV2-SS2 co-infection key proteins. 3.11 Fluorescence experiment of the two pathogens PCV2 and SS2 in PK15 cells As showed in the Figure 10, PCV2 and SS2 were pictured in fluorescence microscope. Meanwhile, gram stain was used for SS2 for comparison with immunofluorescence. This data indicated that PK15 cells were more adherent to bacteria after infected with PCV2 for the amount of SS2 in PCV2 infected cells in larger than PCV2 without cells. Fluorescence experiment verified the two pathogens were executed in all this experiment.

**Figure 9.**
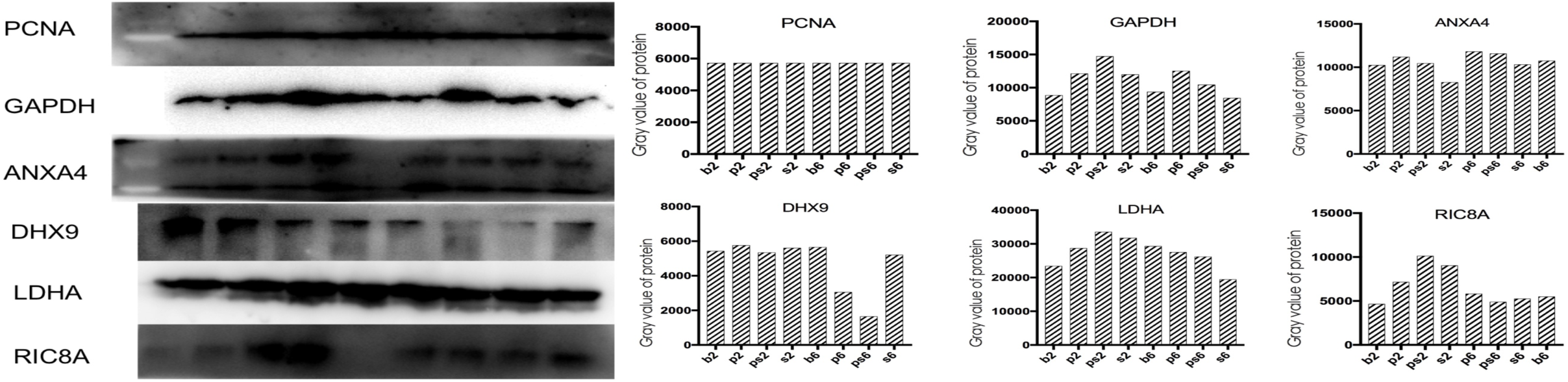
Western blot verification assay. The left of this figure was western blot data of PCNA, GAPDH, ANXA4, DHX9, LDHA and RIC8A, as well as the right of this figure was the statistical data of western blot. The ordinate of the histogram was the gray value of proteins and the abscissa of the histogram were 8 groups.

**Figure 10.**
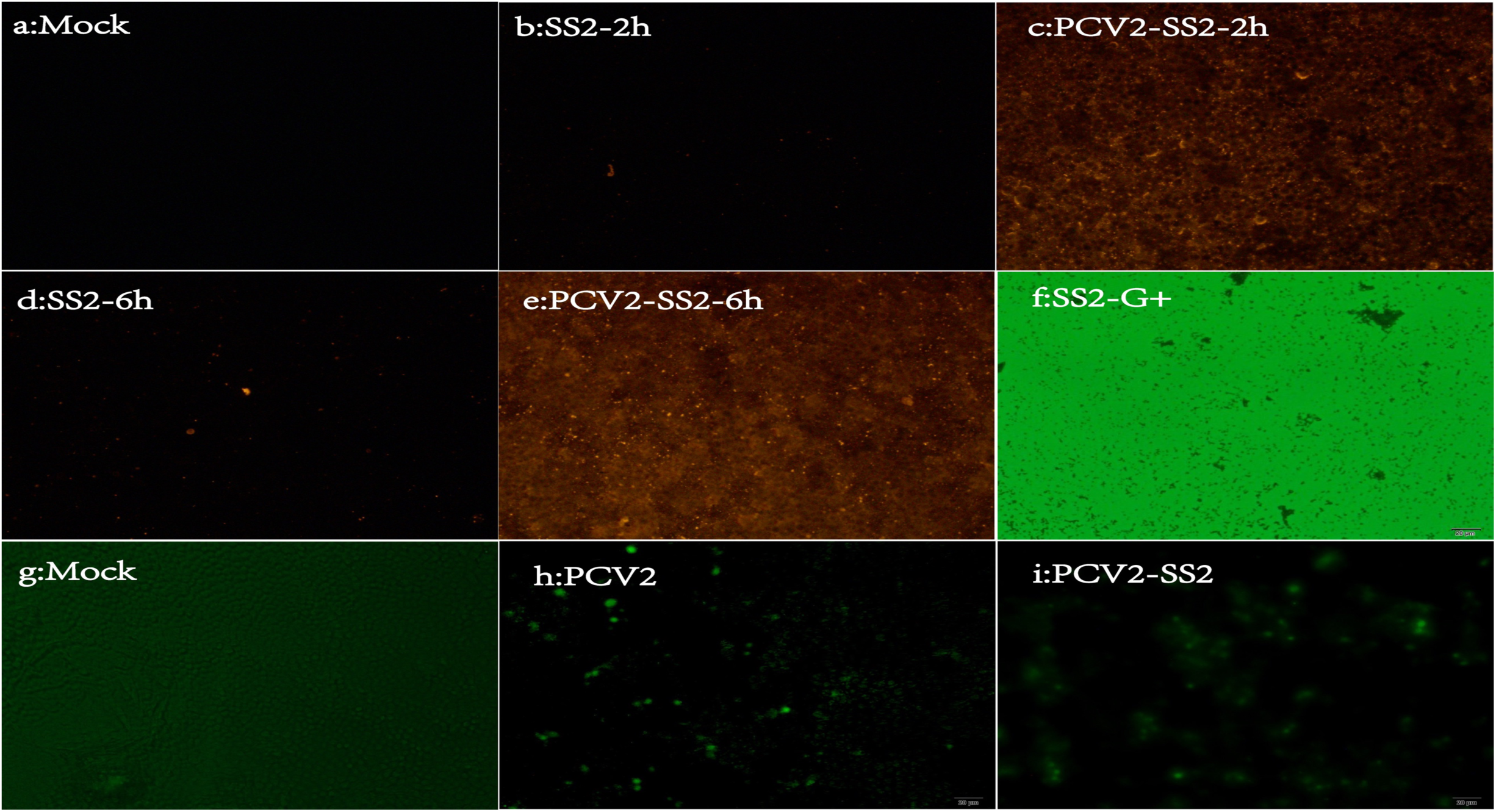
The pictures of SS2 and PCV2 fluorescence stain in fluorescence microscope and Gram stain in microscope. The fluorescence stain of SS2 was marked by CY3 and PCV2 was marked by FITC. Meanwhile, SS2 was also done with Gram stain. a, the blank cells without SS2 infection. b, SS2 infected cells with 2 hours and was showed red fluorescence in fluorescence microscope. c, PCV2 and SS2 co-infected cells with 2 hours and was showed red fluorescence in fluorescence microscope. d, SS2 infected cells with 6 hours and was showed red fluorescence in fluorescence microscope. c, PCV2 and SS2 co-infected cells with 6 hours and was showed red fluorescence in fluorescence microscope. f, SS2 was stained by Gram stain and showed in microscope. g, the blank cells without PCV2 infection. h, PCV2 infected cells and was showed green fluorescence in fluorescence microscope. c, PCV2 and SS2 co-infected cells and was showed green fluorescence in fluorescence microscope. The length of the ruler in the pictures were 20 µm.

## DISCUSSION

When pathogens infect organisms, tissues, and cells, host changes were a complex process, or rather, it was a collection concept. PCV2 could persist existing in this large heterogenous family of myeloid cells (like monocyte, macrophage and dendritic cell) without replicating or being degraded which like a “Trojan horse” ^43^ and PK15 cell was used to culture and isolation of viruses of PCV. PCV2 could infect many cells in vivo, but it caused the slightest lesions in porcine kidney cells, perhaps, it was the response to the infection rather than the infection itself that results in the most damage, death and a general decline into catabolic state to the bystander cells ^2,10,44–46^. PCV2 existing research showed that PCV2 induce apoptosis and autophagy by HSP70, HPS90, ROS, MKRN1, pPirh2 ^47^. In the early infection stage virus could induce pro-inflammatory cytokine expression ^48^. PCV2 is a negative modulates virus in vitro ^49^. Type I and type II interferons induce cellular protein alteration and increases replication of porcine circovirus type 2 in PK15 cells ^50^. PCV2 induces IFN beta expression through increased expression of genes involved in RIG-I and IRF7 signaling pathways ^51^. Study also showed PCV2 lead to thymocyte selection dysregulation ^43^.

Extra interleukin-1 supplementation could help newborn piglets resist the infection of streptococcus suis by the may augment nonspecific immune function ^52^. Studied literature had reported that streptococcus suis has albumin binding activity and plasminogen binding activity and the additional of albumin and plasminogen could increase the virulence of Streptococcus suis to mouse ^53,54^. And some reported that extracellular proteins like cellular fibronectin and collagen I, III and V, fibrin, vitronectin and laminin could bind to Streptococcus suis ^55^. Heat-killed capsular of Streptococcus suis type 2 strains could stimulate tumor necrosis factor alpha and interleukin-6 production by murine macrophages ^56^. It was said that the related diseases caused by Streptococcus suis start from the colonization of bacteria to nasal mucosal epithelial cells, and then follow with either the flow of respiratory tract or the bloodstream ^57^. The mechanisms of bacteria how to be involved in the access that from the bloodstream to the central nervous system were unknown. Simultaneously, it was possible that epithelial cells of the choroid plexus also play an important role in the pathogenesis of the meningitis. The common epithelial cell lines in the laboratory include LLC-PK1, PK (15), A549, HeLa and MDCK, etc. It was reported that SS2 was able to adhere to but not to invade epithelial cells, and the haemolysin produced by S. suis serotype 2 was responsible for a toxic effect observed on epithelial cells, that is to say, it was possible that suilysin positive S. suis strains use adherence and cell injury, as opposed to direct cellular invasion to epithelial cells ^57^. Cell death, caspase activation, and high mobility group box 1 (HMGB1) protein release of porcine choroid plexus epithelial cells during streptococcus suis infection in vitro ^58^. As said above, both pathogens caused significant harm to pigs or even human at clinical. The pity was that predecessor studies were confined to understand the infection and co-infection of the two pathogens from the main thought of a single molecule. We explored the proteomics changes of PK15 cells when co-infected with the two pathogens. As the case of a complex co-infection, experiment was starting from a single cell and using proteomics and bioinformatics to find the differential proteins of different groups, which directly and objectively presented the complex protein changes, suggested possible mechanisms for further study.

Our results demonstrated that PCV2 cause apoptosis for it induce the chaperonin containing tailless complex polypeptide 1 decrease ^59^ and supplemented pathways like Wnt/Ca^2+^, and SS2 could binding to cell extracellular secreted protein ^60,61^ though EMC-receptor interaction. PCV2 caused cell catabolism was greater than anabolic, while SS2 caused a violent cell response. For PCV2-SS2 infection group, in the early stage of SS2 infection, for the before infection of PCV2, the response of cells was compensatory, but in the middle infection of SS2, the toxic protein (of which the effect was similar to amoeba, toxoplasma and carcinogen) was further damage to cells and lead to the decline cells activity. SS2 single infection also caused cell damage, but cell could do some defense not only in a compensatory or passive state.

Simultaneously, the data indicated that PCV2 not only lead to diseases relating to metabolism, but also lead to neurological disease, which may the main reason that PCV2 could cause PWMS and PDNS ^62–64^. In the case that PCV2 made the cells in a passive state, it was more convenient for us to observe the main mechanism of action of Streptococcus suis in PK15 cells, we found that SS2 could decrease the proteins of tight junction, actin cytoskeleton and endoplasmic reticulum which made the cell more dispersive and this derivation could be seen in the figure 13 (The more cells were dispersed, the more SS2 was adsorbed, and promoted each other), which may an evidence that SS2 could pass the blood-brain barrier ^65,66^. And the SS2 migration mechanism in cells was worth to be further study.

Cell survival results indicated that PCV2 increased the cell mortality when SS2 infected (Figure 1). But the proteomics showed that PCV2 decreased the cell response degree when SS2 infected (Figure 9). This may be explained by the infection of PCV2 that left the cells in a state of slaughter by SS2 without defense. This result also inspired curiosity if these two pathogens were infected on immune cells as SS2 could cause a rapid and intense immune response including inflammation response in the early stage. But after all, they are all pathogens.

We listed the proteins that changed in different groups of PK15 cells and made a comparative analysis and fund some common grounds, the two pathogens existed some virulence factor that damage the cells like some pathogenesis of parasites. But the detail proteins of PCV2 and SS2 are don’t know base to our experiment. These common proteins including cell membrane receptor proteins, ribosome-associated proteins, Transport RNA-associated proteins, Lysosome, Purine metabolic, N-Glycan biosynthesis proteins. And we performed an intracellular network interaction analysis about then. In order to facilitate subsequent research, protein pathway analysis was performed on all differential proteins. For reliability, qPCR experiments and Western blot experiments validate our data and immunofluorescence experiments showed our pathogens.

Our experiments revealed that 1 point: proteomics was a good science technology to understand pathogenic infections and pathogenic mechanisms, 2 point: the main reason of PCV2 infection exacerbated the cell survival caused by SS2 infection for PCV2 maked cells in a disintegrated state and the proteins in the cell was reduced and the cell metabolic ability was weakened, so it was in a naive state and lacks defense capabilities, 3 point: except nutritional competition, SS2 can stimulate cells very strongly and produce very strong changes in proteins, which was too harmful to cell.

The changed proteins or mainly pathogenic mechanism of PCV2 were mainly including metabolic pathways, hunting’s disease, lysine degradation, insulin signaling pathway, phagosome, microbial metabolism in diverse environments, endocytosis, oxidative phosphorylation, spliceosome and gastric acid secretion. While, SS2 mainly pathogenic mechanism or changed proteins were including spliceosome, protein processing in endoplasmic reticulum, metabolic pathways, staphylococcus aureus infection, ubiquitin mediated proteolysis, complement and coagulation cascades, aminoacyl-tRNA biosynthesis, malaria, tight junction and regulation of actin cytoskeleton. When PK15 cells were co-infected with the two pathogens and SS2 was the main reason made the cells to death in a short time.

Animals were always exposed to microorganisms or pathogenic environments. This study used PK15 cells as a model to explored a series of protein changes produced by PCV2 and SS2 single and co-infection. Definitely, when pathogens encountered to their host, a complex set of intracellular and extracellular interactions determines the outcome of the infection. Understanding the interactions between those host animal or cell and pathogens was critical to developing treatments and preventive measures against infectious diseases. So, the next study about the special proteins of pathogens-host interactions worth to be further explore.

## CONCLUSION

## Abbreviations

CCK8: Cell counting kit-8
CY3: Cyanine3 carboxylic acid
ECM: Extracellular matrix
FBS: Fetal bovine serum saline
FITC: Fluorescein isothiocyanate
HPLC: High performance liquid chromatography
IDA: Information Dependent Acquisition
IPA: Ingenuity Pathways Analysis
IP-MS: Immunoprecipitation-Mass Spectrometry
iTRAQ: Isobaric tags trace quantification
KEGG: Kyoto Encyclopedia of Genes and Genomes
LC-ESI-MS/MS: Liquid chromatography-electrospray tandem mass spectrometry
MOI: Multiplicity of infection
MSI: Mass spectrometry image
PBS: Phosphate buffer saline
PCAD: Porcine Circovirus Associated Disease
PCV2: Porcine circovirus type 2
PDNS: Swine Dermatitis and Nephrotic Syndrome
PK15: Porcine Kidney cell 15
PWMS: Post-weaning multiple system failure syndrome
SILAC: The stable isotope labeling of amino acids in cell culture
SS2: Streptococcus suis 2
STRING: Search Tool for the Retrieval of Interacting Genes/Proteins
THB: Todd Hewitt Broth
TMT: tandem mass tags
TEAB: triethanolamine buffered

## ACKNOWLEDGEMENTS

This work was supported by the National Natural Sciences Foundation of China (31972679), and Research Project from Jiangsu Key Laboratory for Food Quality and Safety-State Key Laboratory Cultivation Base, Ministry of Science and Technology(2019SY001). The authors would like to express their appreciation to Wuhan GeneCreate Biological Engineering Co., Ltd. for their contributions to the bioinformatics support for this project.

---

**Figure S-1.**
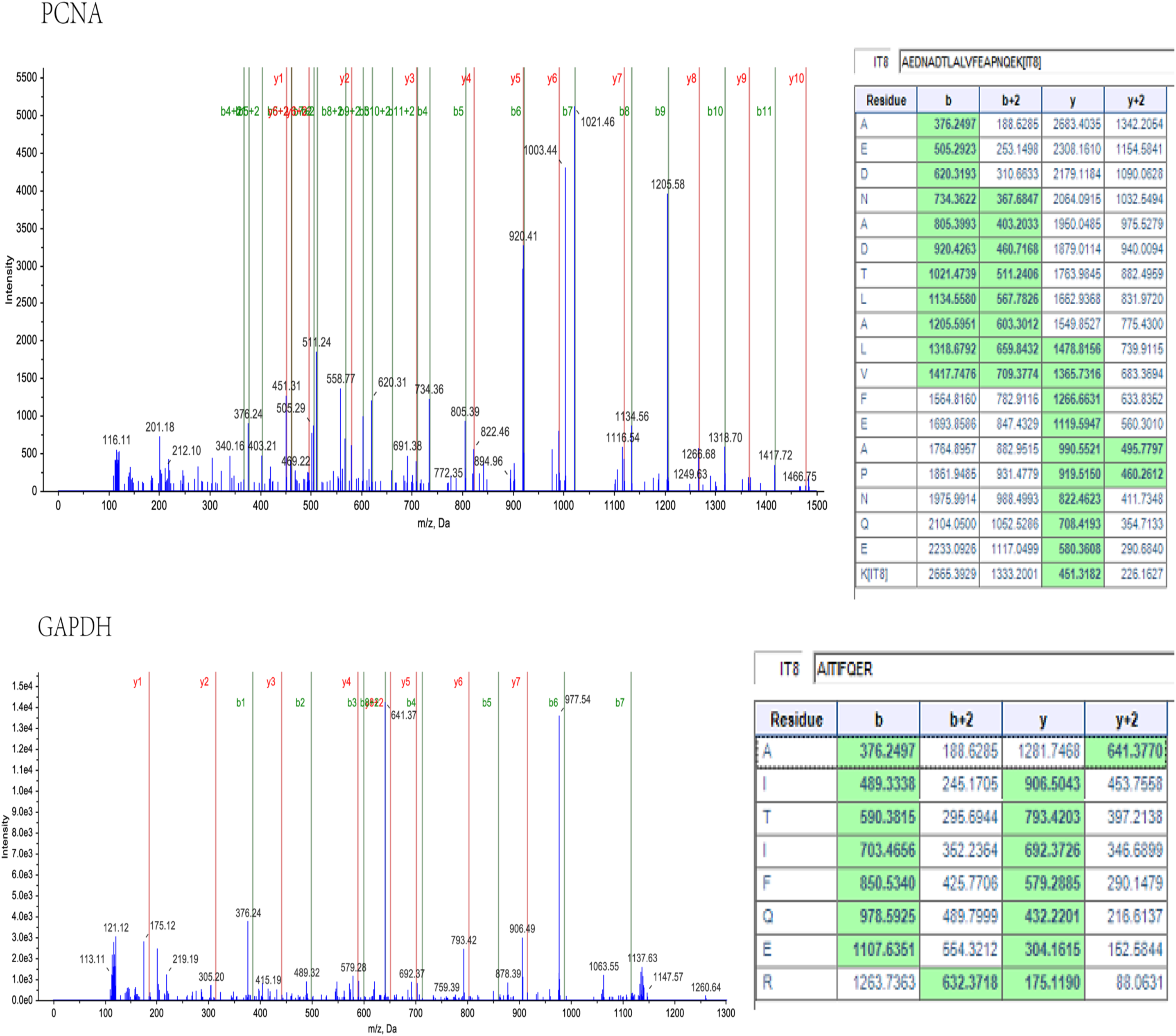
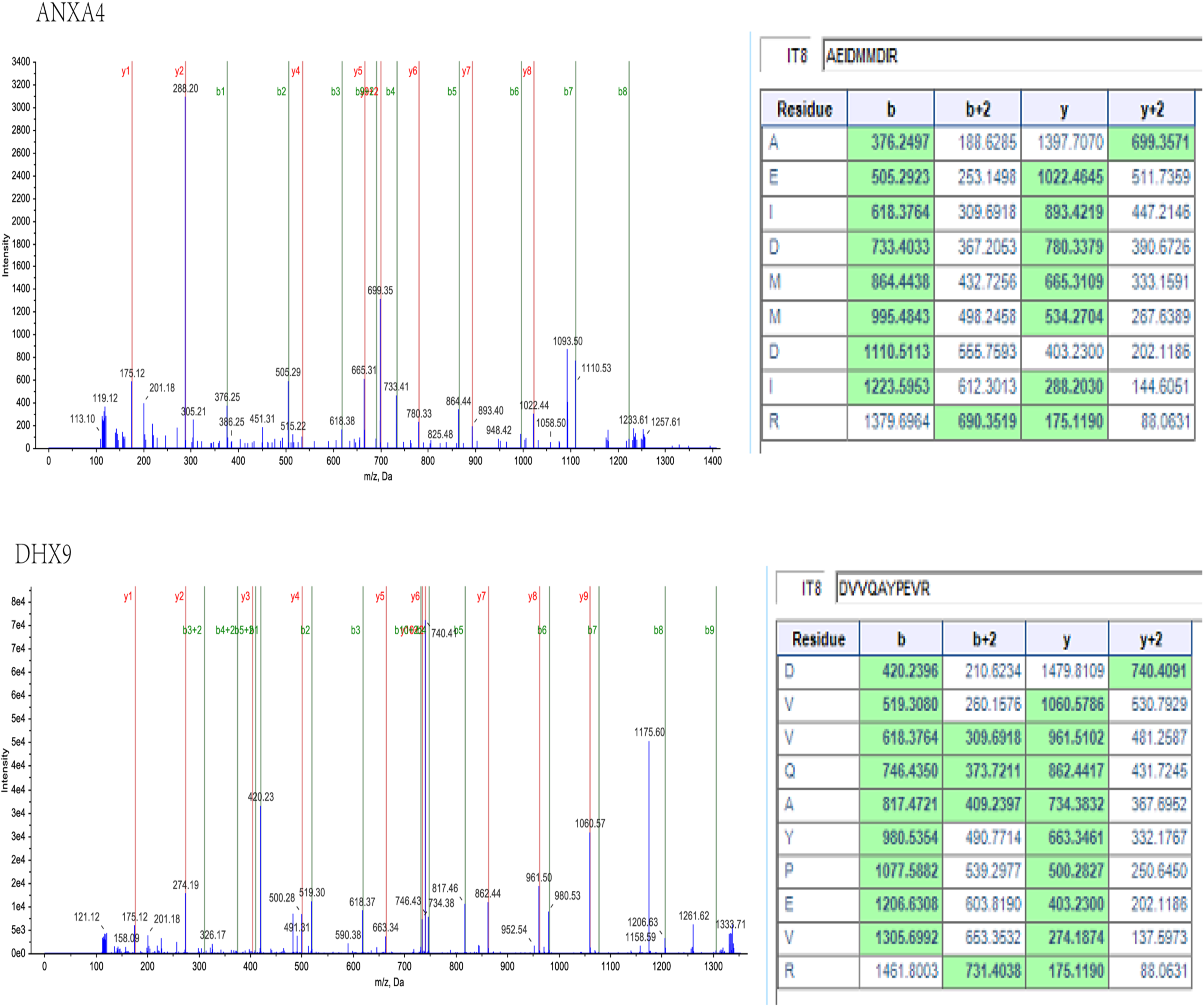
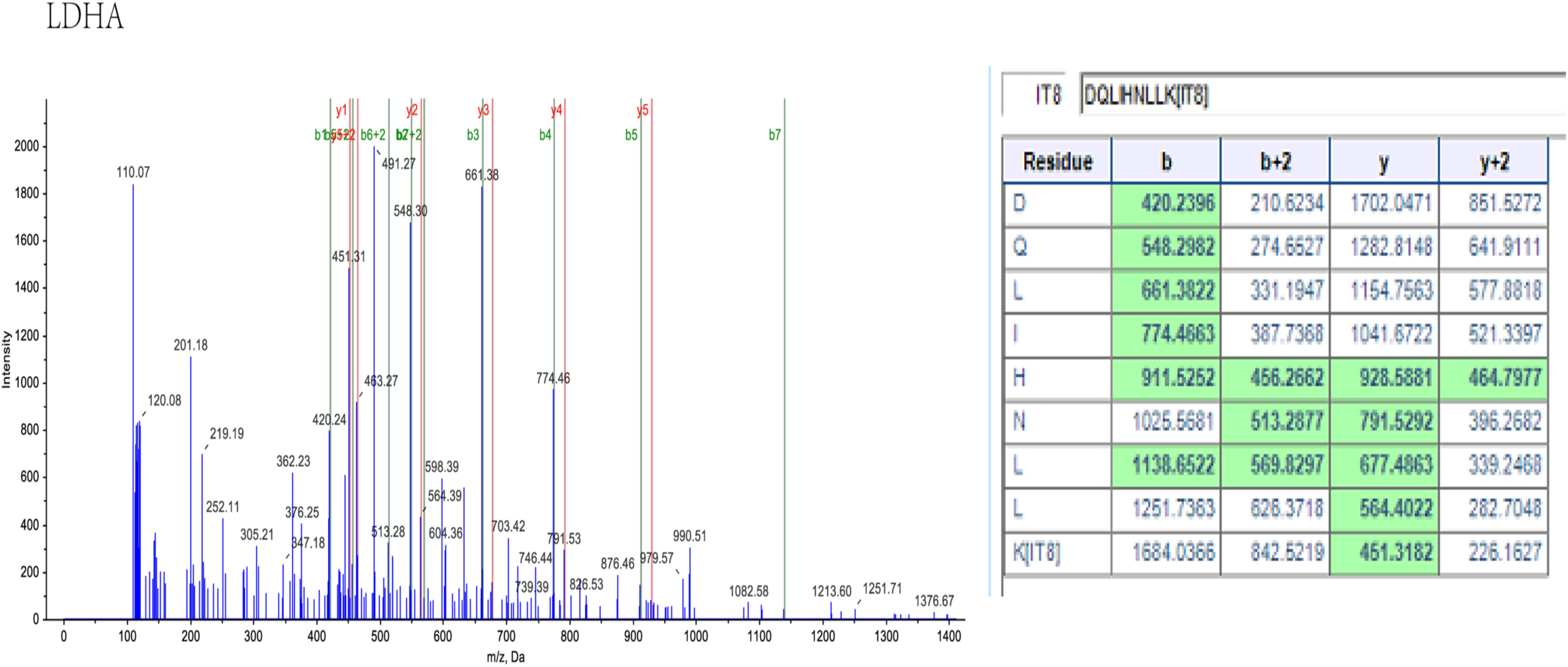
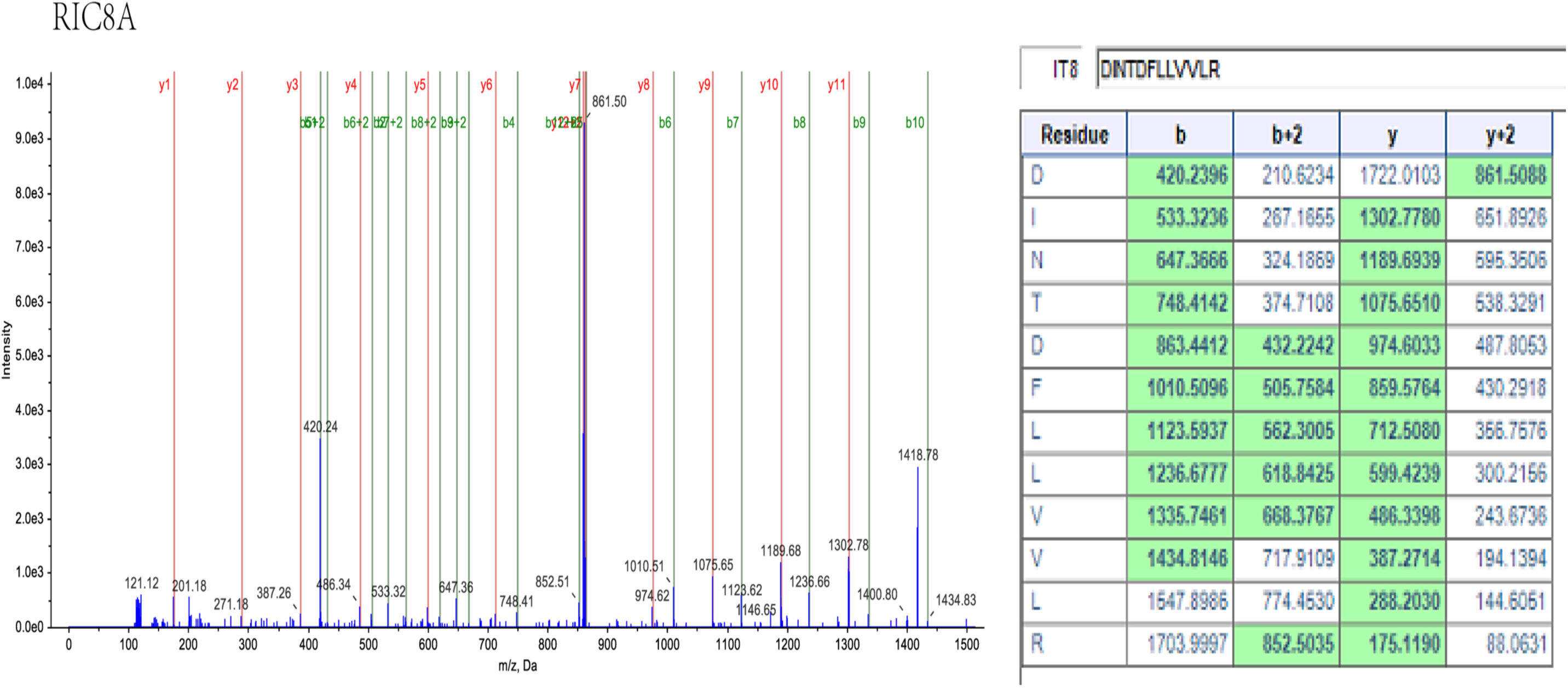
The mass spectrum of protein unique peptide and the mass-to-charge ratio of amino acid in the unique peptide. The mass spectrum figures showed the unique peptide corresponding to a protein, and the tables showed the mass-to-charge ratio of each amino acid in the unique peptide. This figure showed 6 unique peptides of 6 proteins (PCNA, GAPDH, ANXA4, DHX9, LDHA and RIC8A).

**Figure S-2.**
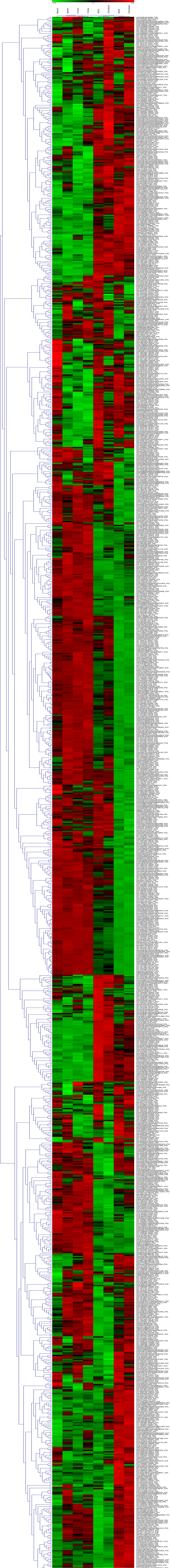
The cluster data of all identified differential proteins in 8 groups. The cluster analysis of identified proteins in 8 groups, of which the ordinate indicated the near and far relationships according unique peptide of different identified proteins and the abscissa indicated proximity of different groups. Green color represented proteins down-regulated, red color represented proteins up-regulated and the color shade degree indicated the degree of up and down regulated. Simultaneously, in the right of the picture was listing the accession number of all these proteins in UniProt database.

